# Intercellular communication controls agonist-induced calcium oscillations independently of gap junctions in smooth muscle cells

**DOI:** 10.1101/734269

**Authors:** S.E. Stasiak, R.R. Jamieson, J. Bouffard, E.J. Cram, H. Parameswaran

**Affiliations:** Department of Bioengineering, Northeastern University, Boston, MA 02115; Department of Biology, Northeastern University, Boston, MA 02115

## Abstract

We report the existence of a unique mode of communication among human smooth muscle cells (SMCs) where they use force to frequency modulate long-range calcium waves. An important consequence of this mechanical signaling is that changes in stiffness of the underlying extracellular matrix can interfere with the frequency modulation of Ca^2+^ waves causing healthy SMCs to falsely perceive a much higher agonist dose than they actually received. This distorted sensing of contractile agonist dose on stiffer matrices is absent in isolated SMCs, even though the isolated cells can sense matrix rigidity. We show that intercellular communication that enables this collective Ca^2+^ response does not involve transport across gap junctions or extracellular diffusion of signaling molecules. The aberrant communication between cells that distorts the individual cell's perception of contractile stimulus can explain the sudden, exaggerated narrowing of the lumen when exposed to low dose of inhaled agonists in diseases like asthma.

## Introduction

Excessive constriction of hollow, tubular transport organs including airways and vasculature is a common pathophysiological feature of widespread disease conditions like asthma and hypertension. The vessel/airway wall undergoes significant pathological changes in the smooth muscle(*1*) and in the extracellular matrix (ECM) that surrounds and supports the cells (*2*) with the onset of disease. The search for mechanisms that underlie the development of these diseases and the search for novel therapies has largely focused on the smooth muscle cells (SMCs)(*3*), as they are the primary effectors of constriction. The pathological changes in the ECM(*2*, *4*), on the other hand, have not received much attention. More recent studies show that changes in the ECM can impact organ function at the very early stages, and can even precede thickening of the muscle layer(*5*). Perhaps, mechanobiological interactions between healthy SMCs and an altered ECM are playing a more critical role in the pathogenesis and progression of diseases like asthma and hypertension than is currently appreciated. In this study, we examined how changes in the matrix stiffness can impact how SMCs sense the dose of an applied agonist.

Agonist induced Ca^2+^ oscillations and long-range Ca^2+^ waves are critical mechanisms that regulate vital parameters such as blood pressure and airway resistance(*6*, *7*). Binding of a muscle agonist, like histamine or acetylcholine, to a surface receptor on the SMC and the subsequent rise in cytosolic Ca^2+^ concentration, is the universal trigger for force generation in the smooth muscle(*8*). Agonist exposure induces Ca^2+^ oscillations in SMCs that propagate as waves within the smooth muscle layer(*9*). These agonist-induced Ca^2+^ oscillations serve two critical functions in the smooth muscle: (A) The concentration/dose of agonist detected by the surface receptors is transduced into the frequency of Ca^2+^ oscillations, with higher concentration of muscle agonist resulting in higher frequency of Ca^2+^ oscillations, which can then be detected by downstream Ca^2+^ sensors and translated into a dose-dependent increase in smooth muscle contractility(*6*, *10*). (B) Agonist-induced Ca^2+^ oscillations propagate as waves around the circumference of the organ and enable the synchronized contractions of SMCs necessary to constrict the airway/blood vessels (*9*). At present, little is known about the role of extracellular mechanical factors such as ECM stiffness in regulating agonist-induced Ca^2+^ oscillations and Ca^2+^ waves that can move across an SMC ensemble.

In this study, we report a collective phenomenon in clusters of human airway smooth muscle, where ECM stiffness alters the intercellular communication between cells in an SMC ensemble causing increased Ca^2+^ oscillation frequencies and synchronized Ca^2+^ oscillations. This altered Ca^2+^ response results in the higher level of agonist-induced force by SMCs on stiff substrates. We examined intercellular transport of Ca^2+^ in SMC cells and we show that contrary to dogma, the physical mechanism that enables intercellular Ca^2+^ waves does not involve molecular transport across gap junctions or paracrine signaling through extracellular diffusion. Rather, this phenomenon appears to be driven by force-transfer among cells in the cluster. The collective response of SMCs to agonist could be a mechanism by which matrix remodeling can drive disease progression in asthma and hypertension.

## Results

### 1. Matrix stiffness alters the Ca^2+^ response to agonist in multicellular clusters of SMCs, but not in isolated cells

#### Modeling the SMC layer in 2D using micropatterning

To study the role of altered matrix stiffness on the frequency of Ca^2+^ oscillations, we used micropatterning to create a 2D approximation of the organization of SMCs seen in lung slices (Fig. 1A). The substrate we use is NuSil(*11*), an optically clear, non-porous polydimethylsiloxane substrate whose Young’s modulus, E, can be varied in the range 0.3 kPa-70 kPa(*11*). Based on measurements of ECM stiffness in healthy human airways, we set the ECM stiffness of healthy human airways to E=0.3 kPa(*12*). This matches the ECM stiffness of small airways (inner diameter<2 mm) which are known to collapse in asthma(*13*). With the onset of airway remodeling, collagen is deposited in the airways and the stiffness of the ECM increases(*2*, *14*). A substrate stiffness of E=13 kPa was used to mimic remodeled ECM. Using a Ca^2+^ sensitive fluorescent dye (Fluo4-AM), we imaged and quantified the time period of Ca^2+^ oscillations in these SM rings plated on soft and stiff ECM. Images were recorded at a rate of 1 per second for 5 minutes following exposure to 10^−5^ M histamine. On soft ECM (E=0.3 kPa), exposure to histamine resulted in Ca^2+^ oscillations with a time period of 44.13 ± 21.76 seconds (N=195, Fig. 1E). When the substrate stiffness was increased to E=13 kPa, the same dose of agonist-induced significantly faster oscillations with a time period of 21.79 ± 8.90 seconds (N=172, p<0.001, Mann-Whitney test, Fig. 1E). Therefore, at the *same* dose of agonist, stiff substrates resulted in a doubling of the cytosolic Ca^2+^ oscillation frequency in healthy smooth muscle cells.

**Fig. 1.**
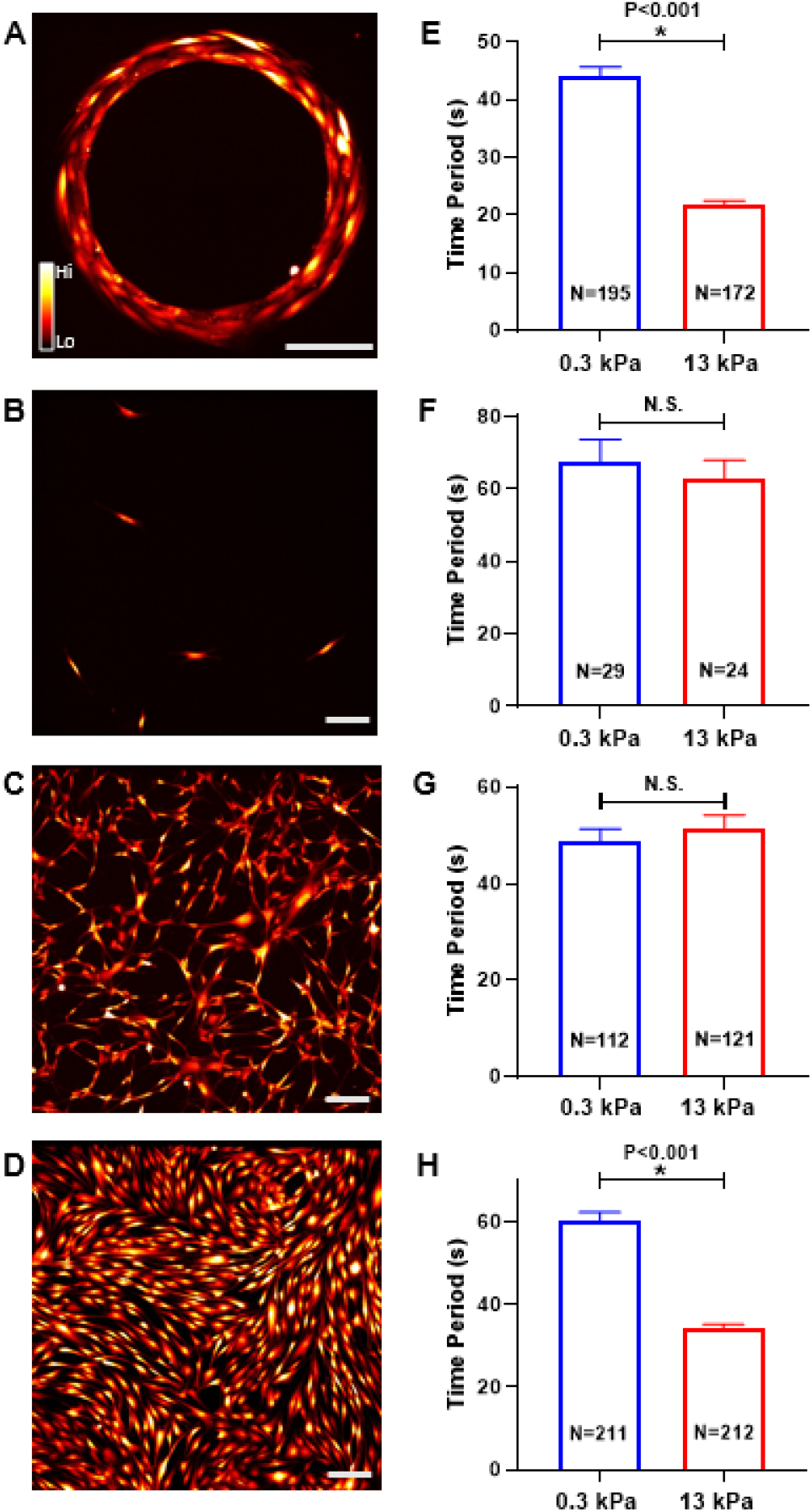
Effect of matrix stiffness on agonist-induced Ca^2+^ oscillations in SMCs. **(A)** To study the role of altered matrix stiffness on the frequency of Ca^2+^ oscillations, we used micropatterning to create a 2D approximation of the in-situ organization of SMCs. Cells were also cultured on non-patterned surfaces in three different seeding densities: **(B)** isolated, **(C)** sparse, and **(D)** confluent. Variations in cytosolic [Ca^2+^] were tracked with a fluorescent Ca^2+^ sensitive dye. Cytosolic Ca^2+^ concentration is pseudo-colored and increases from black to red to yellow to white, following the color bar. Scale bar=200 μm. The mean time periods of agonist-induced Ca^2+^ oscillations were measured for each cell in these conditions on soft (0.3 kPa) and stiff (13 kPa) substrates. **(E)** Increasing matrix stiffness caused a significant decrease in Ca^2+^ oscillation period in SMCs patterned in a ring (Mann-Whitney test). **(F, G)** In both the isolated and sparse culture conditions, SMC Ca^2+^ oscillations were not affected by matrix stiffness, and the mean periods were not statistically significantly different (Mann-Whitney tests, P=0.668, P=0.658, respectively). **(H)** However, in the confluent condition, matrix stiffening caused a large decrease in Ca^2+^ oscillation time period (increase in frequency). The mean period decreased by half from soft to a stiff matrix, as it did in the ring SMCs **(E)** (Mann-Whitney test). The height of each bar represents the mean value across the number of cells, indicated in the figure by N, and the error bar represents the standard error of the measurements. Statistical significance is indicated in figures with an asterisk and labeled p value for the statistical test. Measurements that are not statistically significant are labeled N.S.

#### Interactions between ECM and isolated cells are insufficient to explain altered Ca^2+^ response

To explain the role of matrix stiffness in regulating the Ca^2+^ response to a low dose of agonist, we first hypothesized that this phenomenon was linked to cell-matrix interactions at the level of the individual cell. With increased matrix stiffness, SMCs develop higher cytoskeletal prestress(*15*) opening stretch-activated Ca^2+^ channels(*16*) and potentially increasing the Ca^2+^ flux into the cell. To test this hypothesis, we cultured human airway SMCs at a low density such that individual cells were isolated, spaced at least 100 μm from each other (Fig. 1B). We first measured baseline traction (pre-agonist) stress exerted by these isolated cells on the substrate to confirm that the baseline traction was significantly higher in isolated cells cultured on stiff versus cells cultured on soft substrates (6.98±1.7 Pa, N=16 on soft vs 15.95±5.0 Pa, N=21 on stiff substrate, p<0.001, Mann-Whitney test). We then exposed these isolated SMCs to 10^−5^ M histamine and measured the time period of Ca^2+^ oscillations. Contrary to our expectations, ECM stiffness had no impact on the Ca^2+^ response of isolated SMCs to 10^−5^ M histamine (Fig. 1F). Cells cultured on the stiff substrate had a mean period of 62.65±25.86 seconds (N=24) and those on the soft matrix had a mean period of 67.46±33.79 seconds (N=29), which was not statistically significantly different. Therefore, despite the higher levels of prestress in individual cells, ECM stiffening has no impact on the agonist-induced Ca^2+^ frequency of isolated cells.

#### SMCs sense matrix as a collective and alter their Ca^2+^ response to agonist

To probe this phenomenon further, starting with the isolated SMCs (Fig. 1B), we increased the seeding density (Fig. 1C) until we had a confluent cluster of SMCs (Fig. 1D). At each seeding density, we measured the time period of Ca^2+^ oscillations for SMCs adhering to soft (E=0.3 kPa) and stiff (E=13 kPa) substrates in response to 10^−5^ M histamine (Fig. 1F-1H). Sparsely seeded cells (Fig. 1G) responded similarly to isolated cells (Fig. 1F), exhibiting no statistically significant difference between the agonist-induced Ca^2+^ oscillations on soft and stiff matrix. The time period of oscillations of sparse SMCs on the soft substrate was 48.80±27.33 seconds (N=112) and 51.38±31.22 seconds (N=121) on the stiff substrate (Fig. 1G). Confluent cells behaved like those patterned in a ring, with cells plated on stiffer substrate exhibiting a significantly higher frequency of Ca^2+^ oscillations in response to 10^−5^ M histamine. Confluent SMCs on soft matrix had a mean period of 60.30±29.55 seconds (N=211), and the same healthy, confluent cells on stiff matrix had a mean period of 34.23±13.01 seconds (N=212) (Fig. 1H). These results suggest a collective phenomenon in smooth muscle cells where clusters of confluent SMCs, but not isolated cells, can sense and respond to changes in matrix stiffness. This finding is extremely significant in all smooth muscle pathologies because all downstream Ca^2+^ dependent molecular processes rely on Ca^2+^ oscillations to perceive the external concentration of contractile agonist detected by the G-protein coupled receptors on the cell surface. Here we show that the combination of confluence and ECM stiffness can alter how cells perceive contractile agonist.

### 2. Matrix stiffening synchronizes Ca^2+^ oscillations within a multicellular SMC cluster

We next explored the nature of intercellular communication underlying the collective agonist-induced Ca^2+^ response in SMCs. To do this, we first analyzed the time series of histamine-induced Ca^2+^ oscillations for signs of interactions among the different SMCs within a confluent cluster. After correcting for drift due to photobleaching of the fluorophore, we calculated the cross-correlation coefficient (*ρ*_*i,j*_ ∈ [−1,1]) of the Ca^2+^ oscillations occurring in the *i^th^* cell and the *j^th^* cell in the cluster for all the cells in a cluster. The measured values of *ρ*_*i,j*_ in a typical SMC cluster is depicted in Fig. 2A as a representative 24×24 matrix, with the extreme values *ρ*_*i,j*_ → 1 (pink) indicating that the Ca^2+^ levels in the *i^th^* and *j^th^* cell rise and fall perfectly in sync with each other (perfectly correlated) (Fig. 2B), *ρ*_*i,j*_ = −1 (green) indicating that when the Ca^2+^ levels in *i^th^* cell rises, the Ca^2+^ levels in the *j^th^* cell falls and vice-versa (anti-correlated), *ρ*_*i,j*_ → 0 (white) indicating no correlation in the Ca^2+^ oscillations occurring in *i*^th^ and *j*^th^ cell (Fig. 2C). To avoid the effects of histamine diffusion, the first 60 seconds immediately after application of histamine was not considered in the correlation calculation. Histograms of *ρ*_*i,j*_ measured in isolated SMCs and confluent clusters of SMCs cultured on soft and stiff matrices are shown in Figs. 2D and 2E. Isolated SMCs (Fig. 2D), did not exhibit correlated Ca^2+^ oscillations, with a mean *ρ* of zero, regardless of whether they were cultured on soft (0±0.34, N =435) or stiff (0±0.32, N=276) substrates. However, for cells in a confluent cluster, matrix stiffening caused a statistically significant shift in *ρ* towards positive correlation (Fig. 2E). The mean *ρ* for confluent clusters on soft matrices was 0.09±0.28, N =31125, versus on stiff matrices, *ρ* increased to 0.36±0.31 N =31125. A two-way ANOVA test with confluence and ECM stiffness as independent factors showed a significant interaction between confluence and ECM stiffness (P <0.001). Post-hoc pairwise comparisons with the Tukey test showed a significant difference in the pairwise correlations in SMC clusters due to ECM stiffness, but no difference due to ECM stiffness in the correlations measured in isolated SMCs. There was no systematic trend in the distance between SMCs and the correlation in Ca^2+^ oscillations.

**Fig. 2.**
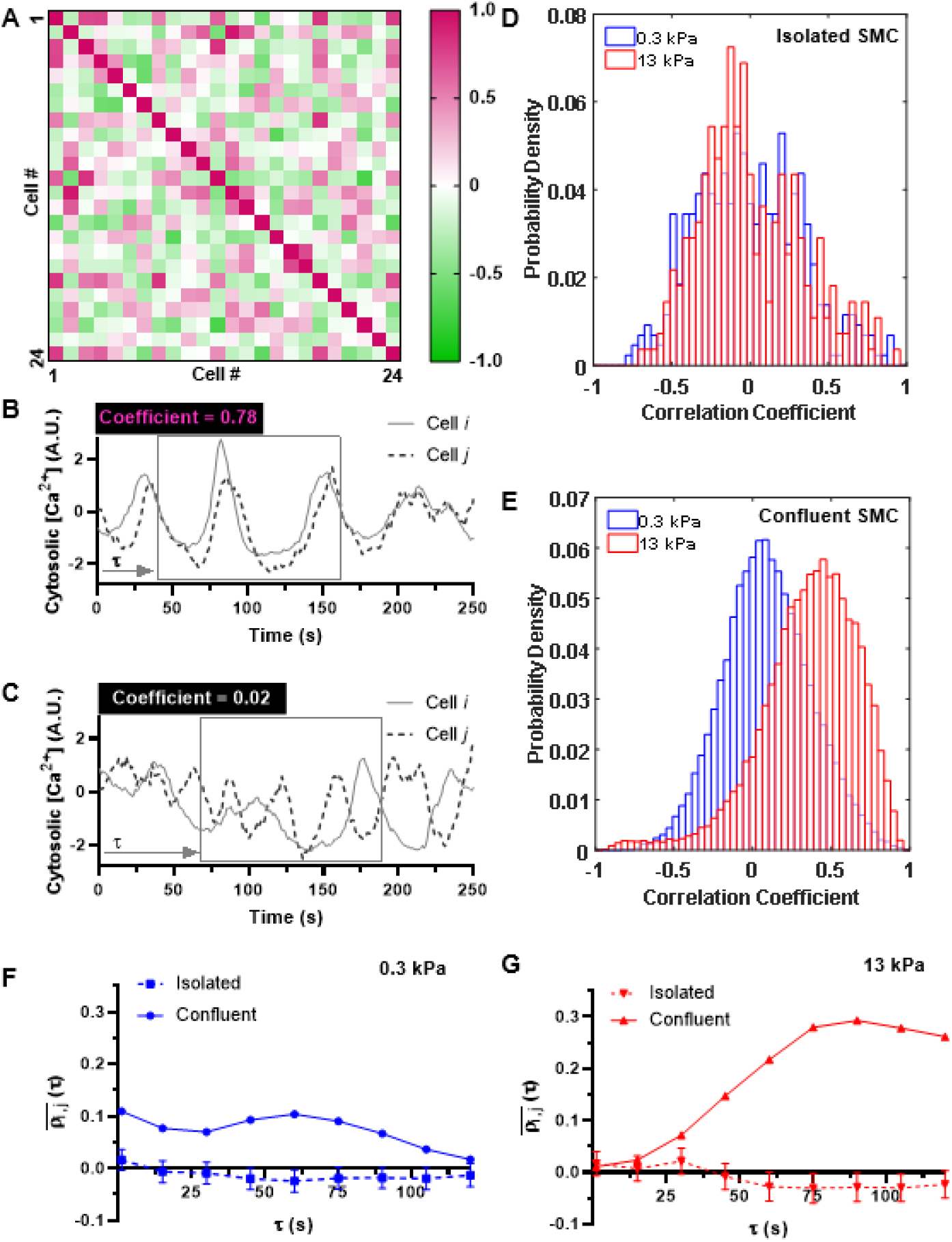
Effect of matrix stiffness and confluence on the correlated nature of Ca^2+^ oscillations. **(A)** The cross-correlation coefficient *ρ* was calculated for the Ca^2+^ oscillation time series of every possible pair of SMCs in confluent and isolated conditions. The coefficients range in values from 1 (positively correlated) (pink) to 0 (uncorrelated) (white) to −1 (negatively correlated) (green), as shown in a representative 24×24 matrix from 24 cells. **(B)** The Ca^2+^ oscillations from a pair of SMCs with a cross-correlation coefficient of 0.78. **(C)** The Ca^2+^ oscillations from a pair of SMCs with a cross-correlation coefficient of 0.02. **(D**) The probability density of correlation coefficients shows that isolated cells on both soft (blue) (N=435 cell-pairs) and stiff (red) (N= 276 cell-pairs) matrix are generally uncorrelated, with the histogram of correlation coefficients centered around 0. **(E)** There is a dramatic positive shift in the correlation coefficient histogram for confluent cells on stiff matrix, indicating that confluent SMCs on a stiff matrix have more synchronized Ca^2+^ oscillations (N=31125 cell-pairs). **(F, G)** To investigate the time it takes after the addition of histamine for the Ca^2+^ oscillations to synchronize, we used a time window of 120 seconds starting at t=60 seconds after addition of histamine and repeated the *ρ*_*i,j*_ calculation for the Ca^2+^ time series within the 120 seconds window for all cells in the cluster. The time window was then shifted in increments of *τ* = 15 seconds over the rest of the 5-minute time frame over which we measured Ca^2+^ oscillations. **(F)** 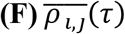 on soft matrix and **(G)** 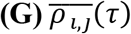 on stiff matrix. Error bars indicate standard error.

Next, we investigated the time it takes after the addition of histamine for the Ca^2+^ oscillations to synchronize. In order to do this, we repeated the previous calculation of cross-correlations, but instead of using the entire time series, we used a time window of 120 seconds starting at t=60 seconds after addition of histamine and repeated the *ρ*_*i,j*_ calculation for the Ca^2+^ time series within this 120 seconds window for all cells in the cluster. The time window was then shifted in increments of 15 seconds over the rest of the 5-minute time frame over which we measured Ca^2+^ oscillations. Shown in Fig. 2F and Fig. 2G is the average through time, of cross-correlation coefficients between all cells either isolated or confluent, 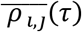 on soft and stiff matrix respectively. The correlation coefficients of confluent SMCs on stiff matrix gradually increase over time, for about 1 minute, which matches the time course for force generation in airway SMCs(*17*). Similar to our findings from the previous section, these results demonstrate that not only does the combination of a stiff matrix and confluent cluster of cells lead to faster Ca^2+^ oscillations, but it also causes histamine-induced Ca^2+^ oscillations to synchronize across the cells in the cluster.

### 3. Gap junctions do not play a role in regulating the collective Ca^2+^ response of SMC clusters

The most well studied long-range communication mechanism in multicellular systems is the intercellular waves of Ca^2+^ capable of propagating over distances much longer than a cell length through a regenerative Ca^2+^ induced Ca^2+^ release mechanism (*18*). This mode of communication is mediated through two critical pathways: (i) gap junction channels, which connect the cytoplasm of neighboring cells and allow for the transport of signaling molecules from one cell to its neighbor and (ii) extracellular diffusion of a signaling molecule like ATP, which can diffuse and bind to purinergic receptors on neighboring cells, causing Ca^2+^ release in these cells (*19*).

#### Transport through gap junctions is unaffected by ECM stiffness

We first fluorescently labeled gap junctions by staining for connexin-43 (Cx43). The expression of Cx43 (red) for confluent clusters of SMCs on soft and stiff substrates are shown in Figs. 3A and 3B, respectively. The actin filaments (green) and the nucleus (blue) are also labeled. From these images, we quantified the number of gap junctions per cell for approximately N=170 cells each for soft and stiff substrate (Fig. 3C). There was no significant difference between Cx43 expression on soft and stiff substrates (Mann-Whitney test, P=0.507). Next, we tested whether ECM stiffness induced a change in the efficiency of transport through gap junctions. To this end, we employed a commonly used technique to quantitatively assess the efficiency of transport through gap junctions called gap-FRAP(*20*). Briefly, a confluent layer of SMCs was incubated with membrane-permeable calcein-AM. Upon entering the cell, the acetyl methyl ester bond was hydrolyzed by intracellular esterases, trapping the hydrophilic calcein molecule within the cell (Fig. 3D, column 1). The calcein in one cell selected at random was then bleached using a high-intensity laser (Fig. 3D, column 2). The bleached calcein molecules and the unbleached calcein molecules in neighboring cells diffuse through gap junctions leading to a recovery in fluorescence (Fig. 3D, column 3). The kinetics and extent of recovery in fluorescence in the bleached cell reflects the efficiency of transport across gap junctions via diffusion. A typical recovery curve from confluent SMCs cultured on soft and stiff matrices are shown in Fig. 3E. We found no difference in the extent of recovery as quantified by the mobile fraction, 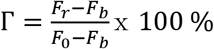, where *F*_0_, *F*_*b*_, *F*_*r*_ indicate the fluorescence intensity in the target cell at baseline, after bleaching and after recovery, respectively (Fig. 3F) (P=0.532, t-test). We also did not observe any statistically significant difference in the rate of recovery in fluorescence *k*_*r*_, which was calculated by fitting an exponential function 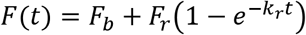 to the recovery curve. On soft ECM, *k*_*r*_ was 0.014±0.018 and on stiff ECM, *k*_*r*_ was 0.011±0.05 (P=0.55, t-test).

**Fig. 3.**
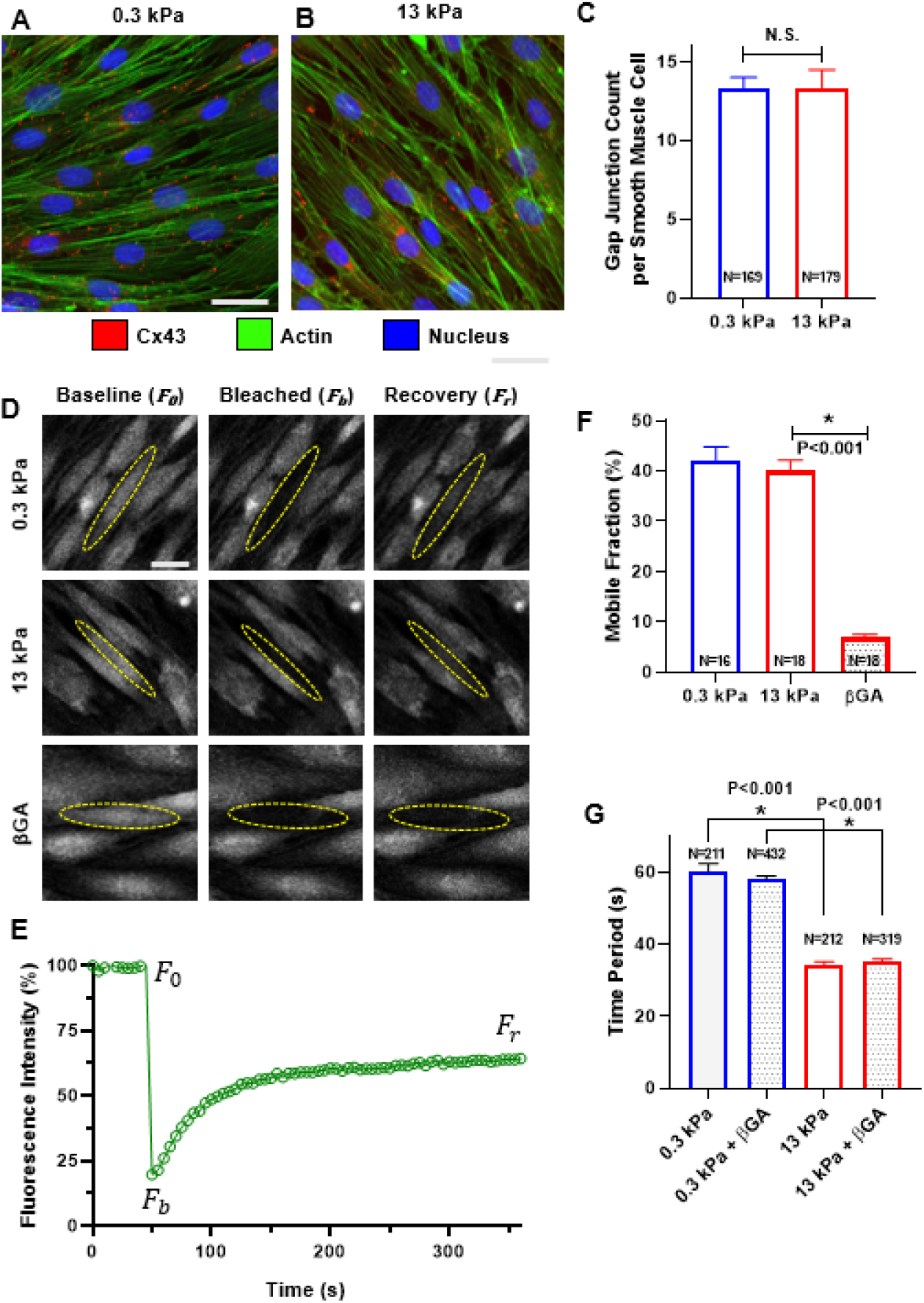
Effect of matrix stiffness on intercellular communication via gap junctions. Confluent SMCs on soft **(A)** and stiff **(B)** matrix was stained for the gap junction protein Cx43 (red), actin (green), and the nucleus (blue). The number of gap junctions per SMC was measured for each matrix stiffness. The mean, number of cells, and standard error are shown in **(C)**. We found no statistical difference between the number of gap junctions per cell on soft vs stiff matrix (Mann-Whitney test, P=0.507). In order to quantify diffusion through gap junctions, we used a technique called gap-FRAP, Briefly, a confluent layer of SMCs was incubated with membrane-permeable calcein-AM. Upon entering the cell, the acetyl methyl ester bond was hydrolyzed by intracellular esterases, trapping the hydrophilic calcein molecule within the cell and it can only diffuse between cells through gap junctions. The baseline fluorescence of a cell was recorded **(D, column 1)**, then the target cell was photobleached to ~20% of its baseline intensity **(D, column 2)**. The fluorescent signal of the bleached cell was recorded for an additional 5 minutes to allow for diffusion of fluorescent molecules between coupled cells, leading to a recovery in fluorescence **(D, column 3)**. **(E)** shows a representative recovery curve over this experiment for one cell which recovered ~40% of its initial fluorescence. These values were used to calculate mobile fraction, a measure of diffusion efficiency through gap junctions. The mean mobile fraction for N cells is shown in **(F)** with standard error bars. Matrix stiffness had no effect on the mobile fraction (P=0.532, t-test). The gap junction blocker 18β-glycyrrhetinic acid (βGA) significantly reduced the recovery **(D, row 3)** and mobile fraction **(F)** (t-test). **(G)** Despite blocking gap junctions with βGA, there was no effect on Ca^2+^ oscillation periods in confluent cells on either soft or stiff matrix (two-way ANOVA). Scale bars= 30 μm.

#### Blocking gap junctions does not affect agonist-induced Ca^2+^ oscillations

To further explore the role of gap junctions in the collective response of the SMCs to agonist, we used 30 μM of 18 β-glycyrrhetinic acid (βGA) to uncouple gap junctions. We first used gap-FRAP to verify that a 30 μM dose of βGA is sufficient to completely block transport across gap junctions. The mobile fraction, Γ, measured in a confluent layer of SMCs after application of 30 μM βGA dropped significantly from 40.05±9.18% to 6.84±2.89% (N=18, t-test, P<0.001) (Fig. 3F). This recovery is similar to the 7.89% recovery we measured in an isolated cell. Similar minimal recovery has been noted in the literature(*21*), indicating that transport across gap junctions was blocked. We then measured histamine-induced Ca^2+^ oscillations in confluent clusters of SMCs. Much to our surprise, we found that blocking gap junctional transport had no impact on the histamine-induced Ca^2+^ oscillation periods in multicellular clusters of SMCs, regardless of the stiffness of the ECM (Fig. 3G). A two-factor ANOVA with treatment (βGA) and ECM stiffness as the two independent factors showed no difference in the time period of histamine-induced Ca^2+^ oscillations (P=0.579) due to treatment. Post-hoc pairwise comparisons showed the Ca^2+^ oscillations remained significantly faster on the stiff matrix (P<0.001) even after blocking gap junctions.

### 4. Mechanical force transfer among cells regulates the collective Ca^2+^ response of SMC clusters

#### Ca^2+^ wave propagation follows the contractile axis of the SMCs

Next, we considered extracellular diffusion of signaling molecules, such as ATP, as a possible mode of intercellular communication that enables the collective Ca^2+^ response of SMC clusters on stiff matrices(*18*). In order to do this, we measured the direction in which the Ca^2+^ wave propagates from one cell to the next in SMC clusters cultured on stiff substrates. If we select a cell in an SMC cluster, a wave that passes through it will appear as a localized increase in Ca^2+^ in the cell which is followed by a localized increase in Ca^2+^ in one of its neighbors. We reasoned that if the intercellular Ca^2+^ transport was being enabled by extracellular diffusion of signaling molecules, then the resulting Ca^2+^ wave should have an equal chance of moving in all directions (isotropic). We split the direction of propagation of the Ca^2+^ wave from an SMC into two directions: a direction parallel to the contractile axis of the SMC, and a direction perpendicular to the contractile axis of the SMC. For the cell labeled 1 shown in Fig. 4A (insets), cells 2 and 4 were considered parallel to cell 1’s contractile axis and cells 3 and 5 were considered perpendicular to cell 1’s contractile axis. We calculated the conditional probability for a localized increase in Ca^2+^ in one cell to be followed by a localized increase in a parallel neighbor versus its perpendicular neighbor. This conditional probability quantifies the isotropy in Ca^2+^ wave propagation with respect to the contractile axis of the SMC, with equal probability (0.5) in parallel and perpendicular direction indicating isotropy in Ca^2+^ wave propagation. Instead, we found that there was an 80.5±7.53% chance for the Ca^2+^ wave to follow the contractile axis of the SMC (Fig. 4B, N =50 cells, Mann-Whitney test, P<0.001). The high probability of the Ca^2+^ wave to follow the contractile axis of the SMCs rules out extracellular diffusion as the dominant mechanism for intercellular communication in our experiments.

**Fig. 4.**
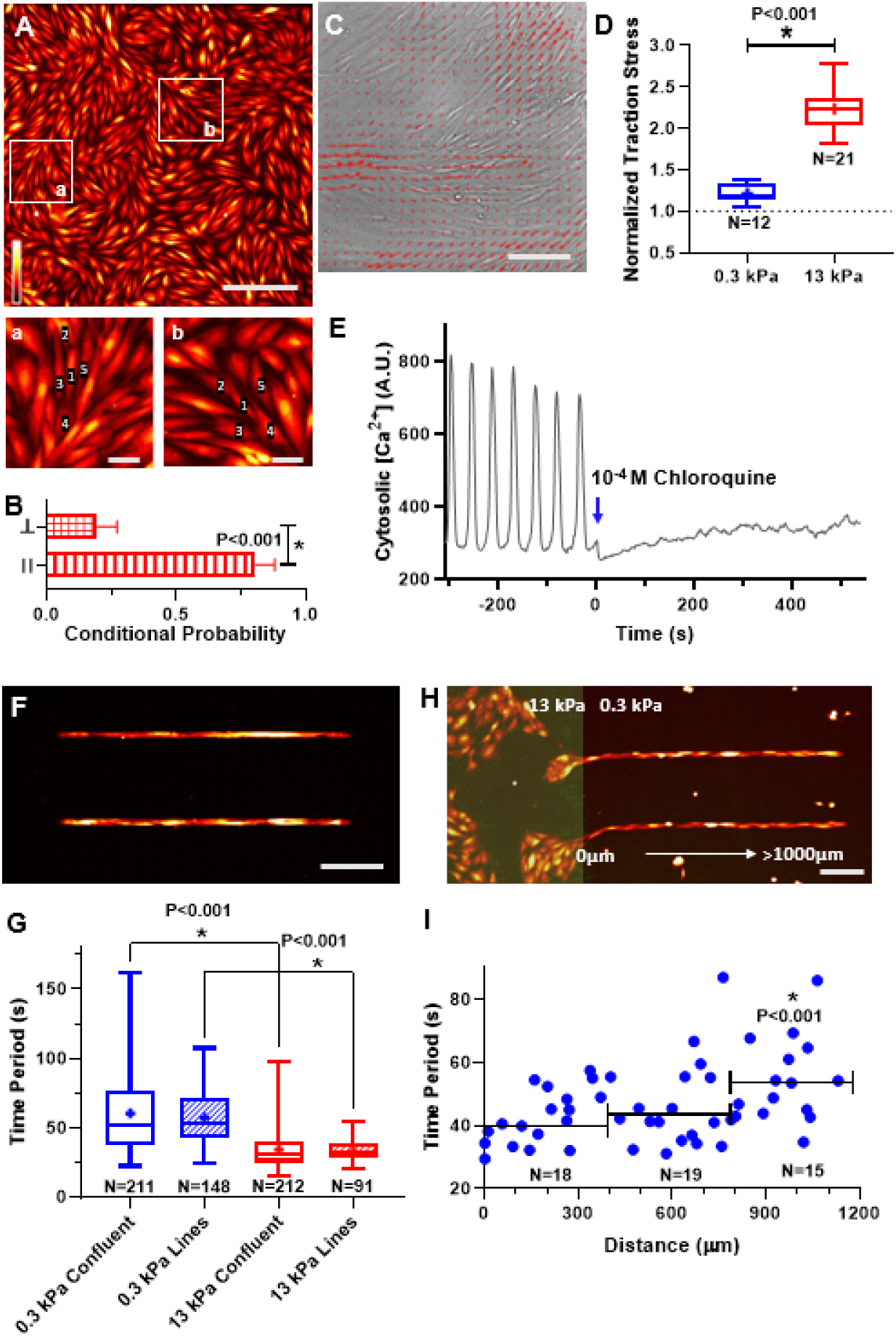
The role of mechanical force in Ca^2+^ wave propagation through multicellular ensembles of SMCs. **(A)** SMCs in confluent layers form organized clusters of cells, with certain cells aligned end-to-end along their contractile axis (parallel), and others branching off at an angle (perpendicular). Scale bar=250 μm. Insets **(a)** & **(b)** show cells 2 and 4 are considered parallel to the contractile axis of cell 1, whereas cells 3 and 5 are considered perpendicular to the contractile axis of cell 1. Inset scale bars=50 μm. The cytosolic [Ca^2+^] is pseudo-colored as in **Fig. 1**. The conditional probability for a localized increase in Ca^2+^ in cell 1 to be followed by an increase in a parallel or perpendicular neighbor is plotted in **(B)** with mean±sd. Ca^2+^ waves were statistically more likely to propagate along the direction of the contractile axis (N=50 cells, Mann-Whitney test). **(C)** Overlaying the traction-stress vectors onto a phase-contrast image of confluent SMCs, we observe that the contractile axis of the cells aligns with the long axis of the cell. Scale bar=150 μm. **(D)** Histamine caused SMC traction to increase by 20% on soft matrix. On stiff matrix, the same dose caused a two-fold increase in force. **(E)** Histamine-induced Ca^2+^ oscillations ceased immediately after exposure to chloroquine, a muscle relaxant. **(F)** When SMCs were patterned in lines, **(G)** the probability of finding SMCs with high time periods decreased and the variance of the time periods from aligned cells was significantly smaller than confluent cells (F-test, P<0.001). Matrix stiffness still affected the oscillation period (Mann-Whitney test). To probe the limits of the SMC cluster’s collective matrix sensing abilities, we simulated localized ECM stiffening by **(H)** patterning SMCs in lines spanning a dual-stiffness PDMS substrate (E=13 kPa highlighted in green, left, and E=0.3 kPa, right). The Ca^2+^ oscillation time period of cells along the lines is plotted as a function of the distance from the stiff matrix **(I)**. Binning cells in 400 μm intervals, only those over 800 μm from stiff matrix had different Ca^2+^ oscillation time periods from cells directly in contact (Mann-Whitney test). **(F)** and **(H)** scale bars=250 μm.

#### Mechanical force transfer between SMCs regulates their Ca^2+^ response

Thus far, we have ruled out transport through gap junctions and extracellular diffusion, the two widely accepted mechanisms responsible for intercellular communication in confluent cell clusters using Ca^2+^ waves(*18*). The propensity of Ca^2+^ waves to follow the direction of the contractile axis of the SMC suggested force transfer between neighboring SMCs as a potential mechanism that regulates the collective Ca^2+^ response of SMC clusters. To quantify SMC forces in our confluent clusters, we used Fourier transform traction force microscopy(*22*) to measure the traction forces generated by 10^−5^ M histamine. Fig. 4C depicts the traction force vectors overlaid on the corresponding confluent SMCs, indicating stress magnitude in vector length and color, and stress direction. We found that histamine generated normalized traction (post-histamine/pre-histamine) of 1.22±0.11, (N=12) on soft substrates. On stiffer matrix, a confluent cluster of SMCs from the same healthy donor and passage generated a normalized force of 2.24±0.26 (N =21) (Fig. 4D). There was a statistically significant difference (P<0.001, t-test) between the force generated on soft and stiff substrates in response to the same dose of agonist. To test the possibility that higher force transfer among cells on stiff substrates was responsible for this collective phenomenon, we tested the effect of reducing the muscle force on the Ca^2+^ oscillations in a confluent cluster of SMCs on stiff substrates. Starting with a confluent SMC cluster which was cultured on a stiff (13 kPa) substrate and exposed to 10^−5^ M histamine for 20 minutes, we measured Ca^2+^ oscillations pre- and post-addition of 10^−4^ M chloroquine, a known smooth muscle relaxant (*23*). Reducing contractility with chloroquine abrogated Ca^2+^ oscillations in the SMC cluster (Fig. 4E). This result suggests contractility is required for the observed collective Ca^2+^ oscillations.

#### Confining the SMCs to a straight line reduces the probability of SMCs with low frequency of Ca^2+^ oscillations

Comparing the alignment of SMCs in Fig. 1B to Fig. 1D and Fig. 4A, we observed that with the onset of confluence, SMCs naturally tend to organize themselves into spatial clusters of aligned cells. To test the effect of SMC alignment on higher frequency of Ca^2+^ oscillation on stiff substrates, we compared the time period of Ca^2+^ oscillations in confluent SMC clusters to the time period of Ca^2+^ oscillations in SMC clusters micropatterned in a line which was (1000 μm long and 15 μm wide, ~1 cell wide and ~10 cell lengths long) (Fig. 4F). The idea here was to eliminate the possibility of intercellular communication occurring perpendicular to the contractile axis in confluent clusters; through mechanisms which are less likely to be influenced by force. We found that while the mean time period of oscillations was nearly identical in both confluent SMC clusters and lines of SMCs, the probability of cells with higher time period of Ca^2+^ oscillations decreases when SMCs are aligned (Fig. 4G). An F-test shows a significant decrease in the variability of the time period distribution when the cells are aligned in a line. (N=115, P<0.001). This result is consistent with the idea that higher Ca^2+^ frequencies are being driven by force transfer along the contractile axis.

### 5. The effect of localized ECM stiffening can be sensed by SMCs over long distances

ECM remodeling in airways and blood vessels often occur as spatially localized processes. How does this pathological change in the ECM spread? Current theories(*24*, *25*) require cells to migrate into the region of stiffer ECM for them to sense the altered matrix and respond by excessive secretion of matrix proteins thereby creating a positive feedback loop that leads to spread of this disease. However, given the collective nature of ECM stiffness sensing in SMC clusters, it may be possible for SMCs located far away from the site of ECM remodeling to detect this localized change even though these SMCs are not physically in contact with the stiff ECM. To test this hypothesis and to quantify the distance over which a localized increase in ECM stiffness would be felt by an ensemble of SMCs, we created a dual-stiffness substrate (Fig. 4H), where the region marked in green has Young’s modulus of 13 kPa and the region in black has a Young’s modulus of 0.3 kPa. We then patterned SMCs in a line starting from the stiff region and extending into the soft region. Cells on soft and stiff ECM were simultaneously exposed to 10^−5^ M histamine, and we measured time period of Ca^2+^ oscillations in SMCs on the soft ECM for 5 minutes. The change in Ca^2+^ oscillation time period was not sudden as one moved from the stiff to the soft ECM (Fig. 4I). Rather, the mean time period increased at a slow rate of 1.48 s/100 μm. We grouped the cells by distance from the edge into 400 μm bins and statistically tested the difference in the time period of Ca^2+^ oscillations between each bin and cells in physical contact with the stiff substrate using the t-test. We found that there was no statistically significant difference between histamine-induced Ca^2+^ oscillations for cells on stiff substrates and cells up to 800 μm (approximately 8 cell lengths) away from the edge (N=20, Mann-Whitney test, P<0.001). This result demonstrates that spatially localized alterations in the ECM can be detected by SMCs far from the site of ECM remodeling, suggesting that ECM pathology can spread through the organ much faster than currently believed.

## Discussion

Increased stiffness of the extracellular matrix (ECM) that surrounds and supports cells in tissue is associated with a number of disease conditions ranging from cancer and fibrosis(*26*) to cardiovascular(*27*), lung(*2*), eye(*28*), and age-related diseases(*29*). Traditionally, pathological alterations in the ECM were thought to be the consequence of disease progression. However, it is now becoming increasingly apparent in many diseases that matrix stiffening precedes disease development and could, therefore, contribute to disease progression(*5*, *30*, *31*). Recognizing the importance of ECM remodeling, clinical trials were undertaken to restore the healthy, homeostatic state of the ECM(*32*). These early efforts were unsuccessful(*33*) and attention has now turned to understanding and targeting the mechanisms by which cells perceive and respond to changes in the ECM(*31*).

Here, we demonstrate a collective phenomenon in smooth muscle cells (SMC) in which matrix stiffness alters the intercellular communication between cells in an ensemble resulting in elevated contractility at low doses of agonist. We show that this collective mechanosensing phenomenon is enabled by crosstalk among cells using Ca^2+^ waves(*18*). A common mechanism of communication involves the molecular transport of Ca^2+^ and inositol trisphosphate (IP_3_) across gap junctions. However, our measurements showed no tendency for molecular transport through gap junctions to differ depending on ECM stiffness. We blocked transport through gap junctions using 18 β-glycyrrhetinic acid (βGA)(*34*) and confirmed that the dose we used was sufficient to completely disrupt molecular transport across gap junctions. Contrary to our expectations based on current models(*18*), blocking transport through gap junctions had no impact on the collective agonist response of SMCs to ECM stiffening. There was no change in the Ca^2+^ oscillation for SMCs on soft and stiff substrates after βGA treatment. Following a recent finding that mechanical forces can synchronize contraction of cardiac myocytes in the developing heart independent of gap junctions(*34*), we tested whether ECM stiffening led to a switch in the mode of communication between cells from molecular signaling through gap junctions to mechanical force based signaling.

Ca^2+^ waves can travel distances far greater than a cell length because Ca^2+^ waves regenerate in each cell by “calcium-induced calcium release” from the endoplasmic reticulum (ER). In the most widely accepted theory of long-range Ca^2+^ transport, Ca^2+^ release from the ER is enabled by a messenger molecule, IP_3_, which diffuses faster than Ca^2+^ and activates the receptors on the ER, so when the Ca^2+^ wave arrives, it can release more Ca^2+^. This theory had its basis in measurements of IP_3_ diffusivity in the Xenopus extract model of the cytoplasm(*35*), which put the diffusivity of IP_3_ at 400 μm^2^/s and that of Ca^2+^ at 40 μm^2^/s. However, more recent measurements(*36*) made in human cells show that IP_3_ diffuses at a slower rate than Ca^2+^. Our finding that molecular diffusion through gap junctions is not necessary to sustain Ca^2+^ waves in SMC clusters is consistent with the challenge to the dogma of Ca^2+^ wave propagation through gap junctions.

In contrast to Ca^2+^ wave propagation in other cell types, measurements in confluent SMC clusters (Fig. 4B) show that Ca^2+^ waves follow the direction of the contractile axis of the SMCs. Further, reducing the SMC force in our experiments with a muscle relaxant stopped the Ca^2+^ oscillations and intercellular Ca^2+^ waves (Fig. 4E) suggesting that mechanical force transfer from one cell to its neighbor enables the intercellular Ca^2+^ transport. Can force transfer between cells also serve as a mechanism which amplifies and modulates the frequency of oscillations as the Ca^2+^wave moves from cell-to-cell? The mechanisms that underlie the findings of the present study can be understood in light of previous work from Felix et al(*37*) and Tanaka et al(*38*) who show that forces applied to a cell membrane cause PIP_2_ to be hydrolyzed resulting in an increase in cytosolic IP_3_ concentration. This would mean that for every SMC cell within a confluent cluster, there are two ways by which IP_3_ can be released into the cytosol: (i) from agonist binding to GPCR receptors on the cell surface. (ii) from IP_3_ generated by a contracting SMC pulling on its neighbor causing IP_3_ release in the neighboring cell(*37*, *38*). This *additional* method of IP_3_ release only exists for cells in a confluent cluster. Further, we have previously shown that force transfer between SMCs increases 8-fold with matrix stiffening(*12*). Such a force-based IP_3_ release mechanism can explain how SMC clusters alter their agonist-induced Ca^2+^ oscillations in a matrix stiffness dependent manner while isolated cells, which lack the second source of IP_3_, do not.

Dysfunction in the smooth muscle has long been thought to be responsible for the exaggerated narrowing of transport organs like as asthma, hypertension and Chron’s disease. Asthma is an example of a disease where remodeling of the ECM is well characterized(*2*), but its effects are not considered in therapy or drug development. Asthmatics can be free of inflammation, have spirometry and respiratory mechanics within the range of healthy individuals up until they are exposed to a smooth muscle agonist at which point, airways in an asthmatic will hyper-constrict (*39*). This exaggerated response of the airway is currently thought to result from sensitization of force-generating pathways in the SMC due to prolonged exposure to inflammatory agents. Here, we demonstrate that the fault may instead lie in changes in ECM stiffness that regulate SMC behavior. We show that a spatially localized change in the ECM might be sufficient for exaggerated force generation in SMCs far away from the site of ECM remodeling. Consequently, the effects of ECM remodeling may manifest much earlier in disease progression than currently believed. Our findings suggest that this exaggerated response to inhaled stimuli might be the result of aberrant communication between cells that distorts the individual cell's perception of contractile stimulus. This is a novel mechanism that has not been explored before in asthma and points to the need to therapeutically target extracellular matrix remodeling for a lasting cure for diseases like asthma.

## Materials and Methods

### Fabrication of optically clear substrates of tunable stiffness

NuSil is an optically clear, biologically inert PDMS substrate with Young’s Modulus tunable in the range from 0.3 kPa to 70 kPa(*11*). Equal parts of NuSil gel-8100 parts A and B (NuSil, Carpinteria, CA, USA) were mixed with various amounts of the crosslinking compound of Sylgard 184 (Dow Corning, Midland, MI, USA) to adjust substrate stiffness. A crosslinker volume 0.36% of the combined parts A and B volume was added for substrates with Young’s modulus E= 13 kPa, and no crosslinker was added to the 1:1 A:B mixture for substrates with Young’s modulus E= 0.3 kPa. After mixing, the substrate was spin coated onto 30 mm diameter, #1.5 glass coverslips for 50 seconds to produce a 100 μm-thick layer. They settled on a level surface at room temperature for 1 hour before curing at 60° C overnight. These cured substrates on coverslips were secured in sterile 40 mm Bioptech dishes (Biological Optical Technologies, Butler, PA, USA) to be used for cell culture.

### Matrix protein coating

In order to coat the entire silicone substrate with protein, a volume of 0.1% gelatin solution was added and incubated at room temperate in the biosafety cabinet for 1 hour. To create protein patterns, we utilized the Alvéole’s PRIMO optical module and the Leonardo software, a UV-based, contactless photopatterning system (Alvéole, Paris, France). The substrate surface was first coated with 500 μg/mL PLL (Sigma Aldrich, St. Louis, MO, USA) for 1 hour at room temperature. The substrate was washed with PBS and 10 mM HEPES buffer adjusted to pH 8.0, and then incubated with 50 mg/mL mPEG-SVA (Laysan Bio, Inc., Arab, AL, USA) at room temperature for 1 hour, and washed with PBS once more. The PRIMO system was calibrated using fluorescent highlighter on an identical substrate. The PBS was replaced by 14.5 mg/mL PLPP (Alvéole, Paris, France), and then the desired pattern, previously created with graphic software, was illuminated with UV light focused on the substrate surface for 30 seconds. Patterned surfaces were washed again with PBS and then incubated with 0.1% gelatin for 1 hour at room temperature. The substrate was washed and maintained hydrated in PBS at 4° C overnight.

### Human airway smooth muscle cell culture

Primary human airway smooth muscle cells (SMCs) were acquired through the Gift of Hope foundation (via Dr. Julian Solway, M.D., University of Chicago) and through ATCC (https://www.atcc.org). Both these sources are public and pertinent medical information about the donor was relayed to us, but all donor identifiers are removed. The donor remains anonymous and cannot be identified directly or through identifiers linked to the subjects, meeting NIH guidelines. This study was carried out in accordance with the guidelines and regulations approved by the Institutional Biosafety Committee at Northeastern University. Cells were grown under standard culture conditions of 37º C and 5% CO_2_ and utilized prior to P7 for traction force experiments and Ca^2+^ imaging experiments. <u>Culture medium:</u> Cells were cultured in 10% fetal bovine serum, DMEM/F12 (Fisher Scientific), 1x penicillin/streptomycin (Fisher Scientific), 1x MEM non-essential amino acid solution (Sigma Aldrich), and 25 μg/L Amphotericin B (Sigma Aldrich). Prior to any measurements, the growth medium was switched to serum-free media for at least 24 hours. The serum-free medium was comprised of Ham’s F-12 media (Sigma Aldrich), 1x penicillin/streptomycin, 50 μg/L Amphotericin B, 1x glutamine (Fisher Scientific), 1.7 mM CaCl_2_ 2H_2_O, Insulin-Transferrin-Selenium Growth Supplement (Corning Life Sciences; Tewksbury, MA), and 12 mM NaOH. Both patterned and non-patterned gelatin-coated substrates were UV sterilized for one hour, then incubated at 37° C for one hour before seeding human airway SMCs, passage 3-6. For patterned substrates, cells were seeded in Bioptech dishes at 10^4^ cells per cm^2^ and incubated for 10 minutes in 10% serum media to allow cells to adhere to patterns. Next, the dishes were washed with PBS to remove excess cells, and then filled with 10% serum media and incubated for 6 to 24 hours. For non-patterned substrates, cells were seeded at the desired density and then incubated in 10% serum media for 6 to 24 hours. After this time, media was replaced with serum-free media and incubated for at least 24 hours prior to measurements. For isolated cells, the seeding density was 10^2^ cells per cm^2^. For sparse cells, the seeding density was 10^3^ cells per cm^2^. For confluent cells, the seeding density was 10^4^ cells per cm^2^.

### Fluorescent imaging of Ca^2+^

Serum-starved airway SMCs were loaded with a fluorescent cytosolic Ca^2+^ indicator to record changes in [Ca^2+^]. FLIPR Ca^2+^ 6 (Molecular Devices, San Jose, CA, USA) was used for all Ca^2+^ measurements except Fig. 1A where we used Fluo4-AM (Sigma Aldrich, St. Louis, MO, USA). The commonly used Fluo4-AM is prone to photobleaching, and the measured Ca^2+^ traces must be bleach corrected prior to measurements of the time period. FLIPR Ca^2+^ 6, on the other hand, did not photobleach even after 30 minutes of continuous imaging at 1 Hz. Fluo4-AM was prepared according to the manufacturer’s standards. Cells were loaded with 0. 2 μM Fluo4-AM solution, diluted in HBSS, and incubated at room temperature for 1 hour. Next, the cells were washed with HBSS and incubated in the dark in HBSS for an additional 30 minutes. The cells were washed once more before imaging. FLIPR Ca^2+^ 6 was also prepared according to manufacturer’s standards. Cells were incubated with a 1:1 solution of FLIPR Ca^2+^ 6 and serum-free media at 37° C and 5% CO_2_ for 2 hours before imaging. Both Ca^2+^ indicators use acetoxymethyl esters to pass through the cell membrane, which are then hydrolyzed by cytosolic esterases, trapping the fluorescent dye inside the cell. Cells were imaged with a Leica DMi8 inverted microscope, a Leica DFC6000 camera (Leica, Wetzlar, Germany), and a Lumencor Sola SEII LED light source (Lumencor, Beaverton, OR, USA). A FITC filter cube (excitation: 480/40 nm, emission: 527/50 nm) was used to image the fluorescent dye. Fluorescent intensity increases with increasing cytosolic [Ca^2+^]. 16-bit images were recorded at 1 Hz for 1 minute before agonist addition, and for at least 5 minutes after 10^−5^ M histamine exposure. In order to analyze data, each image sequence was loaded in Fiji ImageJ, and regions of interest (ROIs) were hand-selected in the cytoplasm of each cell to obtain mean greyscale intensities over the area of the ROI for each frame in time. A custom MATLAB (MathWorks, Natick, MA, USA) code was written to process the data and measure mean Ca^2+^ oscillation periods. This code measured mean Ca^2+^ oscillation periods by finding peaks in the time series data above a certain prominence and taking the mean of the time between all sequential peaks.

### Cell traction force measurements

The base NuSil substrates were coated with a layer of fluorescent beads as fiducial markers for traction force microscopy. A 5% solution of 0.2 μm diameter red fluorescent carboxylate-modified microspheres (FluoSpheres, Invitrogen, Carlsbad, CA, USA) in PBS was vortexed for 10 seconds. 2 mL of solution was added to each substrate in a Bioptech dish and left at room temperature for 1 hour to allow the beads to adhere. The bead solution was poured off, substrates were washed 3x with PBS, and then PBS was poured off. NuSil solution was prepared as described before, with the appropriate amount of crosslinker to match the stiffness of the base substrate. NuSil was spin-coated onto the newly bead-coated base substrate at 2500 RPM to create a 1 μm-thick layer and seal the beads. These substrates rested on a flat surface for 1 hour before curing overnight at 60° C. Substrates were protein-coated and seeded with cells as before. After a 24-hour incubation in serum-free media, the SMC tractions were recorded by imaging the fluorescent beads with a 20x/0.55 dry objective and the Leica DMi8 microscope in an environmental chamber maintained humidified at 37° C. Images were taken at baseline, after 15 minute incubation with 10^−5^ M histamine, and after cells were removed using RLT Lysis Buffer (Qiagen, Hilden, Germany). Using these images, cellular forces were calculated with a custom MATLAB (MathWorks, Natick, MA) software program using Fourier Traction Force Microscopy(*22*).

### Fluorescent labeling of connexin-43/actin/nuclei

Cells were fluorescently labeled for connexin-43 (Cx43) and filamentous actin (F-actin). Cells were fixed in 4% paraformaldehyde in PBS at room temperature for 10 minutes. Then, cells were permeabilized with 100% ethanol for 10 minutes at 20° C. Following permeabilization, cells were blocked using 1x PBS containing 0.1% Tween-20, 1% bovine serum albumin (BSA), and 22.52 mg/ml glycine for 30 minutes at room temperature. Next, cells were stained for Cx43 (ab11370; Abcam, Cambridge, UK) at a dilution of 1:200 in 1x PBS containing 1% BSA for 1 h at 37° C. Secondary antibody labeling and phalloidin staining were done simultaneously at dilutions of 1:200 and 1:40, respectively, using Alexa Fluor 594 (ab150080; Abcam, Cambridge, UK) and Alexa Fluor 488 Phalloidin (A12379; Invitrogen, Carlsbad, CA, USA). Lastly, cells were labeled with NucBlue (Fisher Scientific, Waltham, MA, USA) to label cell nuclei. Images were acquired using a 63x/1.4 oil-immersion objective (Leica, Wetzlar, Germany).

### The gap-FRAP assay

Gap Fluorescent Recovery After Photobleaching (FRAP) is an experimental technique that has been established as an effective method of observing gap-junctional communication of small fluorescent molecules between adjacent cells(*21*). SMCs were cultured on NuSil gels of E=0.3 kPa and E=13 kPa until confluence was achieved, and then serum-starved for at least 24 hours prior to the experiment. Cells were loaded with 1μM calcein-AM solution diluted in warmed 1x PBS solution and incubated at 37° C and 5% CO_2_ for 15 minutes. Calcein-AM is a cell-permeable dye that is hydrolyzed into fluorescent calcein by cytoplasmic esterases upon entry through cell membrane and has been shown to permeate through gap junctions due to its low molecular size (622 Da)(*20*). After dye incubation, samples were washed with warm 1x PBS solution and returned to serum-free medium for experiments. FRAP was performed using a ZEISS confocal laser scanning microscope system equipped with a 20x/0.8 objective and a 488 nm Argon laser. Fluorescence data was captured using ZEN 2012 SP5 imaging software (ZEISS, Oberkochen, Germany). Samples were placed in an incubation chamber maintaining 37° C during experiments to preserve cell viability during imaging. Prior to photobleaching, a manual ROI was drawn around the border of a target cell visibly connected to adjacent cells. Whole cells were selected for photobleaching to ensure that fluorescence recovery could only be attributed to the diffusion of calcein from adjacent cells. Laser power was adjusted to 1% and images were acquired every 5 seconds for 50 seconds using scan speed 12 (pixel dwell = 0.42 μs) to provide baseline fluorescence measurements. After baseline scans were acquired, the laser power was adjusted to 100% and cells were bleached to at least 20% of their initial fluorescence using scan speed 2 (pixel dwell = 40 μs). Following the bleaching step, fluorescence recovery images were collected every 5 seconds for approximately 5 mins at 1% laser power. The images from gap-FRAP experiments were analyzed using Fiji ImageJ software. First, baseline fluorescence intensities from the target cell were averaged over the first 10 frames captured to establish a reference value for fluorescence recovery. Next, fluorescence intensities were measured during the recovery period and divided by the average baseline intensity to normalize the data. To account for any photobleaching occurring during the recovery period, fluorescence intensities were collected from a region at the edge of the field of view. These values were used to adjust the intensity of the target cell over time to account for fluorescence degradation due to repeated scanning of the microscopic field, since the edge region was not affected directly during the bleaching step. The normalized fluorescence values of the bleached cell during recovery were plotted as a function of time. To compare fluorescence recovery across multiple sample groups, the mobile fraction of fluorescent molecules was calculated. Mobile fraction (Γ) measures the fraction of fluorescent molecules that contribute to recovery of fluorescence in bleached cells and is calculated as 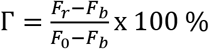, where *F*_0_, *F*_*b*_, *F*_*r*_ indicate the fluorescence intensity in the cell being bleached at baseline, after bleaching and after recovery respectively (Fig. 3E).

### Gap junction blocker experiments

Gap junctions between confluent SMCs were blocked with 18β-glycyrrhetinic acid (βGA) (Sigma Aldrich, St Louis, MO, USA). Confluent SMCs on soft (E=0.3 kPa) and stiff (E=13 kPa) substrates were incubated with 30 μM βGA at 37° C and 5% CO_2_ for 30 minutes. Although βGA is a common gap junction blocker, it has been reported to affect cell viability at higher concentrations(*40*). We used gap-FRAP to find the lowest possible dose of βGA that still blocked gap junctions between confluent cells, which we used here in our experiments. This treatment was used in conjunction with the Ca^2+^ imaging protocol to investigate the role of gap junctional diffusion in agonist-induced Ca^2+^ oscillations.

### Statistical testing

Sigmastat (Systat Software, San Jose, CA) was used to perform statistical tests. Two-factor ANOVAs followed by posthoc pairwise comparisons was used to test for significant differences in datasets which were influenced by two independent factors. Pairwise comparisons used the t-test when the data was normally distributed. Otherwise, the Mann-Whitney test was used to compare the median values. The specific tests used, the number of samples and the p-value are described along with the corresponding results. A p-value of 0.05 was used as the threshold for a statistically significant difference between data sets.

## Author contributions

SS & HP conceived the idea and designed the experiments. With few exceptions, all experimental measurements and data analysis were performed by SS. RR and JB performed the gap-FRAP experiments. RR also performed the gap junction staining. SS, RR, JB, EJC & HP contributed to writing the manuscript and analysis of the data. HP is the corresponding author who conceived and directed this project.

## Data and materials availability

Data supporting the findings of this study are available within the manuscript. All other relevant data are available from authors upon reasonable request.

